# VQ-MAP: a flexible, versatile framework for animal behavior understanding

**DOI:** 10.64898/2025.12.04.692429

**Authors:** Tianqing Li, Ugne Klibaite, Jumana Akoad, Joshua H. Wu, Timothy W. Dunn

## Abstract

The brain generates behaviors that reflect configurations and computations of neural systems. Animal pose analysis can comprehensively profile behaviors in terms of stereotyped action expression, but the absence of common tracking standards makes it difficult to relate measurements across labs and experiments, hindering data exchange and collective knowledge building. Here, we present VQ-MAP, a versatile approach capable of mapping 2D or 3D keypoint sequences with divergent pose formats into a common set of discrete behavioral motifs. VQ-MAP utilizes permutation-invariant attention blocks for keypoint-agnostic processing and a quantized latent space for fast, interpretable delineation of motifs organized into type-subtype hierarchies. We deployed VQ-MAP to align and harmonize heterogeneous pose datasets, co-profiling rodent behaviors in multiple species and models of autism. In longitudinal recordings of rat behavioral maturation, VQ-MAP revealed developmental trajectories connecting specific behaviors across periods of substantial bodily growth and change. By unifying heterogeneous behavioral measurements, our framework will accelerate neurobiological discovery and help the community exercise the full potential of large-scale behavioral datasets.

## Introduction

Understanding the neural underpinnings of animal behavior requires quantification and delineation of the animal behavioral repertoire, along with measurements of behavioral responses to experimental perturbations^1–5^. Videography-based 2D and 3D pose tracking methods now support large-scale measurement of full-body movements in freely behaving animals across a wide range of species, experimental contexts and laboratories^6–14^ (**Fig. 1a**). The expanded scale and resolution of these data establish new lines of behavioral inquiry while presenting new challenges^4,15,16^: how can informative behavioral descriptions be derived from raw kinematic measurements while being reproducible across subjects and robust to heterogeneous data sources across labs and setups?

**Figure 1:**
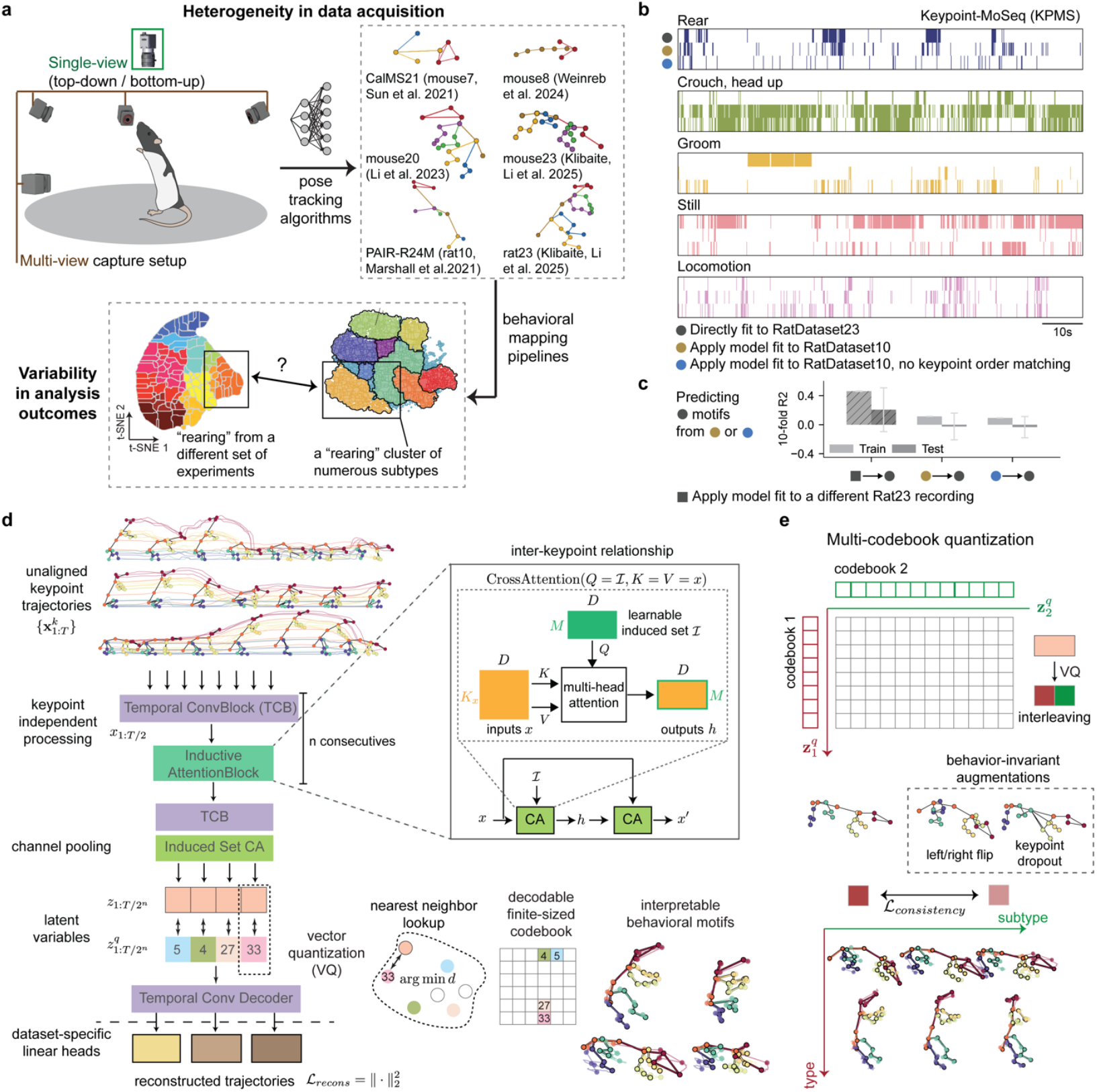
VQ-MAP supports collective processing of heterogeneous behavioral data. **a**, Sources of variability present in the current animal behavioral data acquisition and analysis pipelines, hindering comparative analyses and collective knowledge building. **b**, Raster plots of behaviors identified by Keypoint-MoSeq (KPMS) in an example sequence from RatDataset23, which uses the rat23 3D pose configuration shown in **a**, with the KPMS model directly fit to the indicated datasets. When applying the RatDataset10-fit KPMS model to RatDataset23, we used the 10-keypoint subset common to both datasets, with (middle raster rows, yellow dot) and without (bottom raster rows, blue dot) matching the data structure ordering of keypoints between KPMS training (on RatDataset10) and testing (on the RatDataset23 10-keypoint subset). **c**, Quantification of consistency between the motif identification results in **b**, using linear regression to map between different models. **d**, Schematic illustrating the VQ-MAP modeling framework. Encoder of VQ-MAP embeds and downsamples the pose sequences via *n* consecutive layers of temporal convolutional block (TCB) and inductive attention blocks that capture inter-keypoint relationships via cross attention (CA). Index numbers for vector quantization are only for illustration. **e**, Illustration of the multi-codebook vector quantization architecture for capturing type-subtype hierarchy. Prior to quantization, latent features are partitioned along the channel dimension and independently quantized by separate codebooks. Behavior-invariant data transformations (dashed box) and latent consistency constraints give rise to behavior type-subtype disentanglement.

Recent unsupervised behavioral analysis methods build off of the assumption that naturalistic behavior can be decomposed into series of stereotyped, recurring action motifs (e.g., grooming, rearing, locomotion)^3,17–19^ that are hierarchically organized into finer-grained subtypes (e.g., low and high rearing)^20–22^. These methods quantitatively profile behavior in terms of motif expression frequencies and transition probabilities and, on longer timescales, recurring behavioral sequences and states^23–25^. Stereotyped behaviors can be identified in postural time series by (1) fitting a dynamical statistical model, such as autoregressive hidden Markov models (AR-HMM)^19,26^, which jointly optimize for the motif transition probability and pose dynamics, or (2) clustering of manually crafted sets of postural and kinematic features^18,21,25,27^. While these methods have already driven important neurobehavioral insights^28–31^, several technical issues are throttling their broader impact.

Motif identification remains inefficient and its interpretation challenging, owing to the unstructured output of analyses. Existing methods do not explicitly model the relational hierarchy among motifs (e.g., “turning” → left/right turning subtypes), necessitating secondary post hoc partitioning or subjective human annotation and grouping (**Fig. 1a, bottom**). Manual grouping processes are particularly laborious for methods capturing finer-scale kinematics, as these methods can identify hundreds of distinct motifs^20^. Automated *post hoc* strategies^21,32^ use secondary statistics, instead of kinematic features, as hierarchical determinants, such as assuming behavioral subtypes share the same motif transition statistics^21^.

Current approaches also do not accommodate measurement heterogeneity across labs and setups. Kinematics datasets vary with the identity and number of tracked keypoints and with individual annotation styles (**Fig. 1a, top right**). There exists no keypoint annotation schema sufficiently standardized among annotators and laboratories, and it is impractical to expect any one schema to capture consistent quantities across different species. Without strategies for co-analyzing a range of keypoint configurations, we cannot conduct collective analyses of behavioral kinematics data collected under different experimental setups, strains and species. Moreover, overfitting to specific body plan choices, which may not guarantee to provide a representative abstraction of the underlying behavioral dynamics, can inject unwanted biases (e.g., keypoint order, connectivity) into the behavioral repertoire and limits the transferability of motif identification into new contexts^16^.

Here, we introduce VQ-MAP, a framework enabling integrative, transferrable behavioral analysis by decoupling extraneous dataset characteristics from underlying body movements. VQ-MAP is a variational autoencoder (VAE) yielding discrete vector-quantized (VQ) latent variables^33^, an unsupervised architecture developed to learn a vocabulary of modality units (e.g., image tokens, speech phonemes) combined with autoregressive models for efficient generative modeling^34–39^. VQ-MAP encodes 2D or 3D pose time series as a sequence of discrete candidates drawn from multiple finite-sized ‘codebooks’. The codebooks capture behavioral motifs that are automatically organized hierarchically, with one codebook identifying high-level behavioral categories (rearing, locomotion, etc.) and the other codebook capturing low-level postural and kinematic variations to identify motif subtypes. To accommodate different pose formats, VQ-MAP extracts temporal features from each keypoint time series independently, then uses an attention mechanism to account for cross-keypoint relationships with invariance to input channel cardinality and permutation^40,41^.

Using tracked 2D and 3D poses of freely moving rodents (mice and rats), we show that the channel-independence property of our encoder allows for co-profiling of heterogeneous datasets with variable keypoint cardinality and arrangement, event across different species, and zero-shot encoding of unseen input formats without retraining. In summary, the proposed framework presents multi-pronged solutions for processing of variable inputs and extrapolating the learned representations to novel applications.

## Results

Unsupervised animal behavior profiling approaches segment tracked pose sequences into discrete action motifs^19,20^. However, individual experiments and datasets use ad hoc pose definitions that vary in keypoint cardinality (how many keypoints), indexing (how keypoints are arranged within the defined skeletal structure), and anatomical localization (where keypoints are annotated on the body). To test whether these differences impact combined behavioral analyses across pose configurations, we fit Keypoint-MoSeq (KPMS)^26^ models to two rat datasets acquired with different hardware: RatDataset23, a 23-keypoint 3D pose format (rat23) tracked from multi-view videos using s-DANNCE^42^, and RatDataset10, a 10-keypoint 3D pose format (rat10) tracked using marker-based motion capture (PAIR-R24M^43^). Grossly, rat10 keypoints are a subset of those in rat23, but the precise keypoint locations differ slightly for the body parts common to both datasets.

While KPMS models fit separately to each dataset were able to identify typical human-identifiable motifs (e.g., rearing, locomotion, **Supp. Fig. 1**), other than via qualitative human annotation there was no way to relate identified motifs across models. When we tried applying the model fit to RatDataset10 to RatDataset23, using their common 10-keypoint subset, the resulting motif patterns were disrupted, showing frequent deviations from the motifs identified when fitting directly to RatDataset23 (**Fig. 1b,c, Supp. Fig. 1**). Motif deterioration was even more dramatic when we did not explicitly match keypoint order across the datasets (**Fig. 1b**). These problems were not solved by jointly fitting to both datasets; when co-fitting, identified motifs were segregated with little overlap between datasets (**Supp. Fig. 1e**,**f**). Similar generalization issues were more exaggerated between two 3D mouse datasets whose keypoint sets did not completely overlap (MouseDataset14 vs. MouseDataset20, **Supp. Fig. 1c**,**d, Supp. Table 1**).

Differences in body plans and pose annotation schemas limit our ability to identify behaviors and reason over a diverse range of datasets. Manually aligning keypoints from different pose schemas does not retain model performance (**Fig. 1**) and is not always feasible. Furthermore, existing methods do not learn explicit and decodable representations of postural dynamics^44^, which restricts how the identified behavioral motifs can be quantitatively examined and extended across contexts. When pose formats differ, generalization to new data requires referencing the existing repertoire, for example, by co-embedding with previous data samples or subjective manual motif alignment (that is, the experimenter decides Motif A from Repertoire X represents the same type of rearing as Motif B from Repertoire Y). To reliably and flexibly support quantitative comparisons across behavioral datasets, we need embeddings that explicitly encode kinematic information while being agnostic to as many format-specific details as possible.

### VQ-MAP enables flexible processing of animal postural data

To overcome the challenge of keypoint registration across datasets, we developed VQ-MAP, an unsupervised neural network framework for behavioral motif identification with keypoint format invariance that emphasizes underlying kinematics (**Fig. 1d**). VQ-MAP takes postural time series in arbitrary keypoint formats as input and learns a shared behavioral embedding space unifying these heterogeneous measurements. We based VQ-MAP on the hypothesis that different pose formats represent different “views” of the same underlying behavioral motifs that could be aligned by learning format-specific transformations across multiple datasets. This idea resembles a fundamental approach in computer graphics, “motion retargeting”, where a common animated motion can be transferred between articulated figures with diverse body plans^45,46^.

VQ-MAP separately encodes individual keypoint trajectories using temporal convolutions, then applies a Transformer attention module^40^ to reason about relationships between keypoint temporal features in arbitrary pose schemas (**Fig. 1d)**. After temporal compression, VQ-MAP transforms these continuous kinematics into a stream of discrete, generalized units that facilitate downstream analysis, modeling, and interpretability. Specifically, input 2D or 3D pose sequences are parsed into a series of latent variables, which are then quantized by mapping them to entries in finite-sized codebooks (“vocabulary”), where each entry is characterized by a learned code vector. Unquantized latent variables are mapped, based on their distance to each code vector, to the nearest codebook entry. Like autoencoders, VQ-MAP is trained to reconstruct the original motion sequences, with linear decoders attached to the output accommodating different pose formats if multiple datasets are being used for training. Except for these linear decoders, the entire network architecture is shared across different pose formats, as the core motivation is to decouple dataset characteristics from the abstraction of underlying body kinematics. Decoders are only used for model training; once fit, VQ-MAP does not need new decoders to code-map new pose formats.

When testing VQ-MAP with a single latent codebook, we found that while the model successfully identified behaviors, the learned codes were, like with existing methods, unorganized, with subtypes of the same behavior represented by disjoint codes (**Supp. Fig. 2a**,**b**). To encourage the latent space to organize hierarchically, we imposed additional constraints during training^47,48^. Instead of learning a single, flattened embedding space, we explicitly distribute the latent factors over two codebooks. During training, we applied augmentations to input pose clips (left/right mirroring and random keypoint dropouts, which did not alter motif type) and added a term to the training objective to enforce augmentation-invariance in one of the codebooks (Codebook 1). This self-supervised training strategy produced data-driven type-subtype hierarchical representations in the quantized latent space, where each motif was represented as a combination of entries drawn from both codebooks (**Fig. 1e, Supp. Video 1**). Codebook 1 (CB1) comprised high-level codes capturing common and distinct behavioral categories while codebook 2 (CB2) captured low-level information describing kinematic variations (e.g., vigor, heading direction) under high-level codes (**Supp. Fig. 2c**). For example, code 1 and 5 in CB1 represented rearing and locomotion behaviors respectively. In CB2, code 14 captured, with generality, motifs where the body turns to the right. So, the code pair (1, 14) corresponded to instances where animals twisted to the right during rearing and (5, 14) to right turns while walking (**Fig. 1e**).

**Figure 2:**
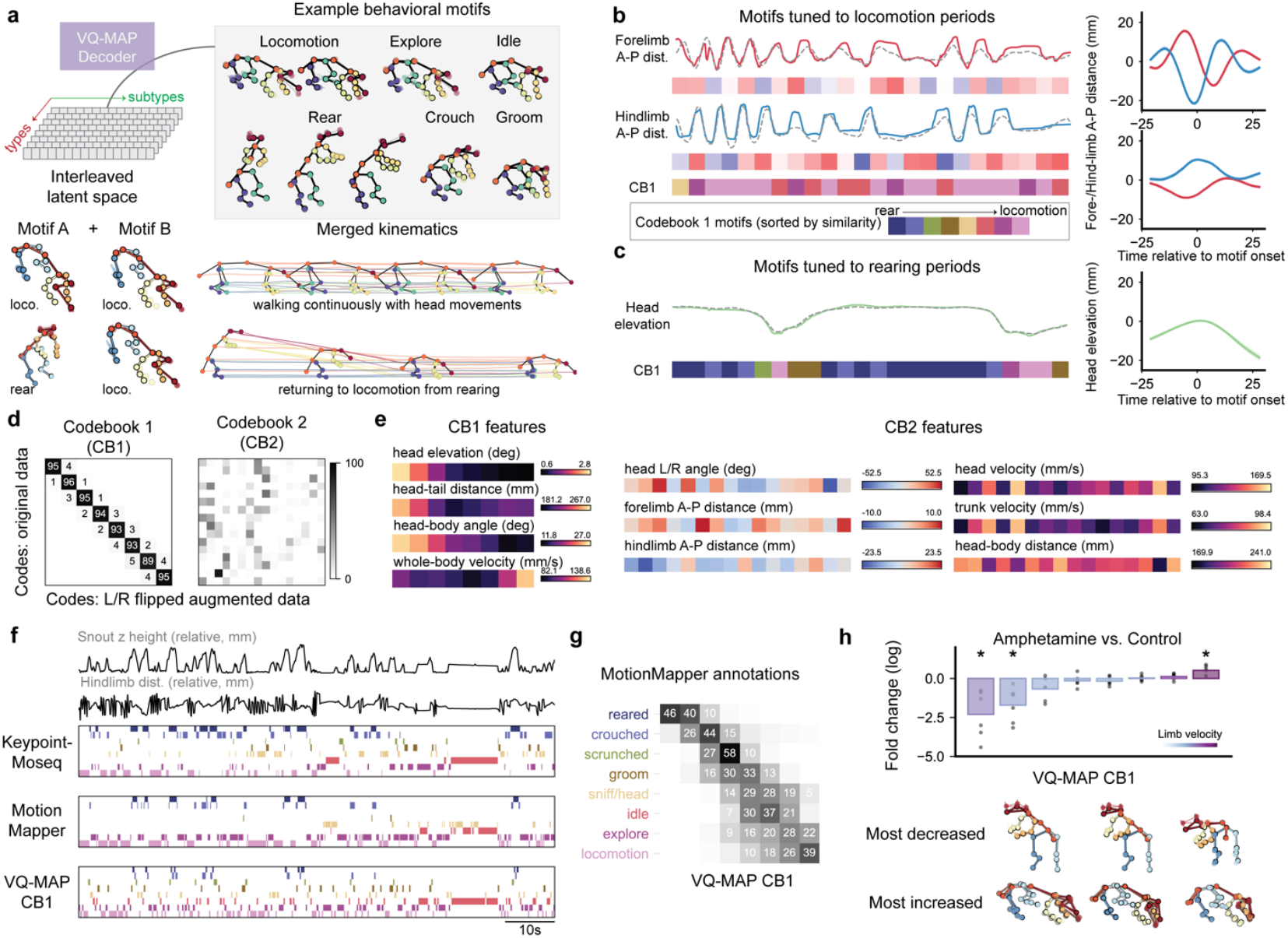
VQ-MAP recapitulates standard behavioral analyses and establishes motif hierarchy without post hoc processing. We fit a VQ-MAP model to RatDataset23 (n = 25 30-minute recordings, n = 5 rats). **a**, Top, example VQ-MAP behavioral motifs from directly decoding the code vectors into kinematics. Bottom, demonstration of how different codes are chained and merged into coherent kinematics by the decoder. **b**, Left, oscillatory forelimb and hindlimb movements (anterior-posterior, A-P, distance between left and right limbs) and corresponding CB1 code mappings in an example locomotion sequence. Solid and dashed lines show A-P distances computed from original pose inputs and VQ-MAP reconstructions, respectively. The cool-warm heatmaps represent the relative A-P distances in the decoded code kinematics. Right, limb movement trajectories for pose sequences mapped to the same VQ-MAP codes (that is, motifs). Two different codes are displayed. Solid lines represent the mean, shaded regions 95% CIs. Codebook 1 (“CB1”) code vectors were sorted by descending cosine similarity relative to the code representing the highest amplitude rearing motif for easy visualization. **c**, Left, example rearing period and corresponding CB1 codes. Right, mean ± 95% CIs head trajectories for a rearing-specific code. **d**, Confusion matrices showing code mapping consistency on a held-out set of recordings (n = 5) with left/right lateral flipping transformations. **e**, Mean kinematic and postural features calculated from pose sequences directly decoded from motifs. **f**, Behavioral ethograms from Keypoint-MoSeq, MotionMapper with human annotations, and VQ-MAP CB1 codes, for an example sequence. Each row of an ethogram corresponds to a different type of behavior and shows when each behavior was expressed (color legend in **b**). Corresponding snout height (a proxy for rearing) and hindlimb movements (a proxy for locomotion) are shown for reference. **g**, Confusion matrix showing overlap between human labeled behaviors (from MotionMapper cluster annotations) and VQ-MAP CB1 categorization. The value in a cell, reflected also in grayscale level, is the percentage of samples of a given MotionMapper type assigned to a given VQ-MAP CB1 code (rows sum to 100). **h**, Top, fold change in VQ-MAP CB1 code usage before and after amphetamine administration (RatAmphDataset, n = 6). Bottom, decoded VQ-MAP motifs for codes with the largest usage changes (* p < 0.05 t-test).

### VQ-MAP recapitulates standard analyses and captures behavioral hierarchy

To validate VQ-MAP, we asked whether it could recapitulate standard behavioral analyses. We started by embedding RatDataset23 using codebook sizes of 8(CB1):16(CB2) and characterizing the postural and kinematic attributes associated with the learned behavioral motifs. The decoder reconstructs individual codes into fixed-length body movement bouts (**Fig. 2a, top**) and joins longer chains of codes into coherent kinematic sequences (**Fig. 2a, bottom**). Via manual inspection, we identified codes corresponding to common rodent behavioral types (locomotion, grooming, rearing, idling, etc., **Supp. Fig. 2**). Via decoding, we could also directly visualize and analyze the kinematics represented by each code. For instance, within those representing locomotion, we found codes differentially attuned to initiation step laterality and gait frequency (**Fig. 2b**). Some codes were specifically attuned to different stages of rearing, with associated CB1 motifs consistently mapping to changes in head elevation (**Fig. 2c**).

We next investigated the representations of type-subtype hierarchy supported by kinematic feature disentanglement in CB1 vs. CB2 (**Supp. Fig. 1, Supp. Fig. 2c**). To quantitatively benchmark disentanglement, we embedded a held-out test set of recordings and, separately, a version of the test set with left/right flipping augmentations applied. Corresponding sequences were consistently mapped into the same CB1 codes but diverged across CB2 (**Fig. 2d**). We further examined the features represented in each codebook from decoding motif kinematics. CB1 codes were mostly distinguishable by the general postural parameters, such as body elevation (‘head z height’) and elongation (‘head-tail distance’), and CB2, as expected, demonstrated a more diverse range of subtle movement patterns with pronounced differences in body laterality, limb oscillatory movements and body part-specific velocities (**Fig. 2e, Supp. Fig. 3**).

**Figure 3:**
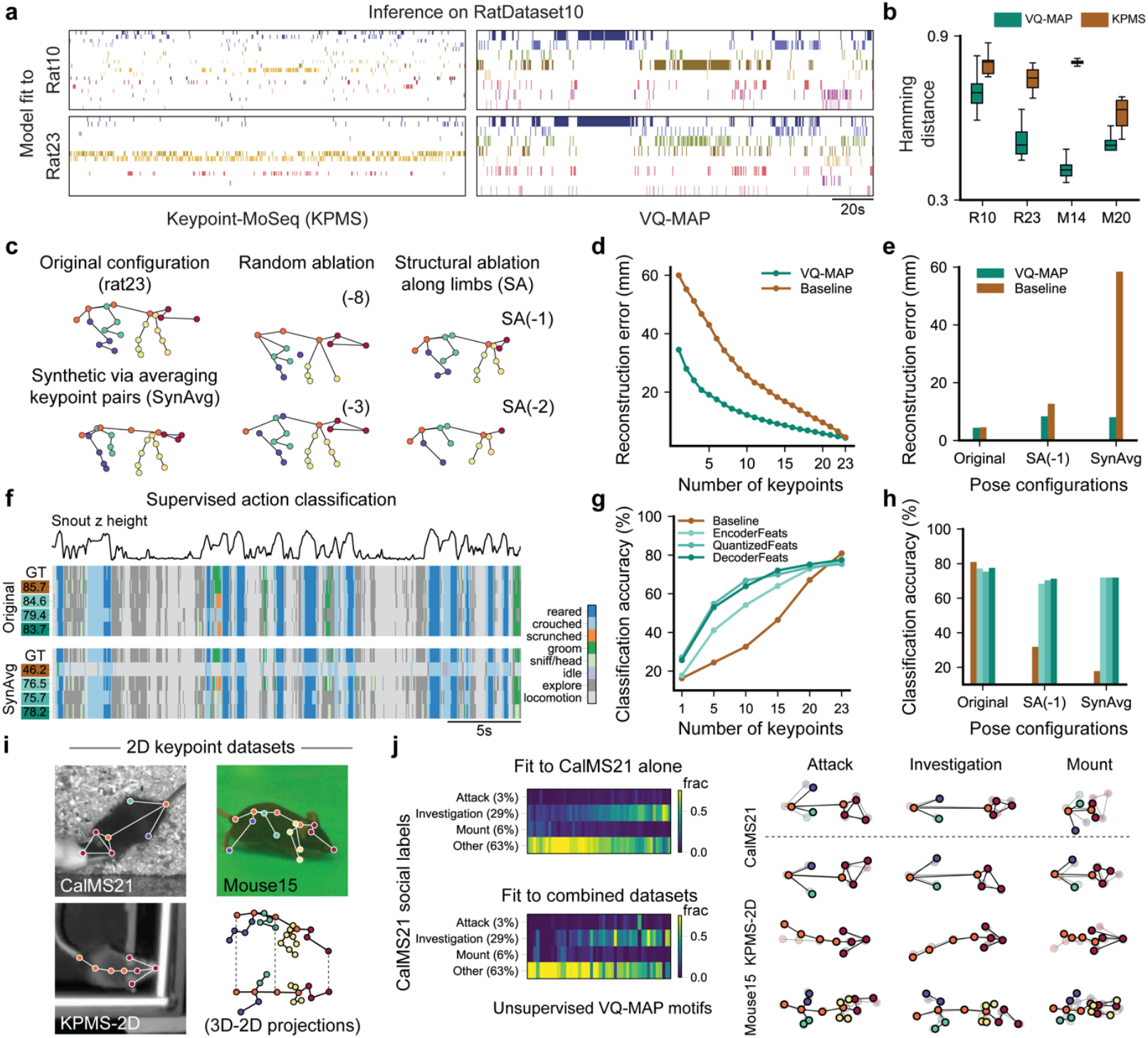
VQ-MAP identifies shared behavior across variable datasets. **a**, Comparison of behavioral ethograms for an example RatDataset23 recording (cf. **Fig. 1b**) between KPMS (left) and VQ-MAP (right). Each row represents a different motif, colors are used here only for visual contrast. KPMS motifs were manually sorted from rearing to locomotion to be consistent with VQ-MAP. **b**, Quantification of motif consistency between different model outputs (**Methods**). **c**, Schematic of skeletal ablations for benchmarking the robustness to noise and perturbations applied to the rat23 3D pose configuration. **d**, Mean pose sequence reconstruction error as a function of input keypoints, assessed using random ablations. The baseline model (brown) is a VQ-VAE lacking channel-invariant processing. **e**, Comparison of reconstruction error between original rat23 and the indicated pose perturbations. **f**, Comparison of action classification results on an example recording with original rat23 (top) and SynAvg (bottom) poses, across four LSTM classifiers trained using the original poses (“Baseline”, brown) and intermediate features drawn from VQ-MAP model’s encoder (“EncoderFeats”, light aqua), latent bottleneck (“QuantizedFeats”, mid aqua) and decoder (“DecoderFeats”, dark aqua). GT, ground truth. **g**, Quantification of classification accuracy as a function of tracked keypoints, assessed using random ablations. **h**, Comparison of classification accuracy between using original rat23 and the indicated pose perturbations. **i**, Example frames from the 2D mouse datasets included in 2D VQ-MAP model training. **j**, Left, confusion matrices showing overlap between CalMS21 human-annotated social behaviors and VQ-MAP unsupervised motifs in the resident animal, for a model fit only to CalMS21 (top) and to the combined 2D datasets (bottom). Values in columns sum to 1, that is each value in a column is the fraction of VQ-MAP codes that overlap with a human-annotated CalMS21 social behavioral label. Right, VQ-MAP motifs most associated with each of the indicated CalMS21 behavioral categories, for the CalMS21-only and combined dataset models.

To assess the fidelity of the learned type-subtype categorization for behavioral motif mapping, we directly compared the ‘ethograms’ for RatDataset23, which were directly produced using inferred CB1 mappings without post hoc annotation and grouping, with those produced by two common motif identification approaches, Keypoint-MoSeq (KPMS)^26^ and MotionMapper (MM)^18,25,29^ (**Fig. 2f, Methods**). Motifs produced by MM were annotated and grouped into 8 general behavioral categories by two human labelers^42^. Despite the variability in modeling formulations, motifs found by all methods shared highly overlapping semantics, i.e., they qualitatively described the same core behaviors (**Supp. Fig. 1**). The behavioral segmentation results produced by VQ-MAP revealed substantial similarities to the two existing approaches, both qualitatively and quantitatively (**Fig. 2f**). Similar to KPMS and MM, three VQ-MAP CB1 codes captured different types of rearing behaviors and the two CB1 codes different forms of animal locomotion. Further comparison of VQ-MAP CB1 codes with human-annotated MM motif clusters revealed that CB1 codes corresponded with human-recognized behavioral categories (**Fig. 2g**). While CB1 code alignment with human-defined categories aids interpretability, we note that such alignment is not strictly necessary for quantitative comparisons across experiments.

Last, we assessed VQ-MAP’s ability to detect behavioral changes in response to experimental perturbations. In data from n = 6 Long-Evans rats with and without amphetamine injection (**Methods, Supp. Table 1**), VQ-MAP revealed amphetamine-associated modulation of high-level CB1 codes for rearing and locomotion, agreeing with previous MM reports^42^ (**Fig. 2h**).

### VQ-MAP identifies shared behaviors across datasets within species

VQ-MAP was designed to reconcile diverse data formats in a single model, both during training and when extrapolated to unseen pose schemas. To evaluate the framework’s flexibility in this regard, we first assessed the cross-dataset motif identification performance (cf., **Fig. 1b,c**). VQ-MAP mitigated the issues observed for KPMS when testing on datasets with different keypoint cardinality from the training source (e.g., RatDataset10 model → RatDataset23 and vice versa) (**Fig. 3a, Supp. Fig. 1a**). VQ-MAP cross-dataset motifs closely matched with those identified by models directly fit to the target rat dataset. To quantify cross-dataset motif identification performance, we fit linear models to predict cross-dataset from within-dataset motifs to account for differences in motif ordering and, for KPMS, in the number of identified motifs (10-fold cross validation, mean ± 95% CI Hamming distances between held-out test motifs and model predictions, target dataset: VQ-MAP vs. KPMS, Rat10: 0.70 ± 0.05 | 0.76 ± 0.07, Rat23: 0.49 ± 0.07 | 0.73 ± 0.05, **Fig. 3b, Supp. Fig. 1a**,**b**). This trend was exacerbated in mouse datasets (Mouse14: 0.42 ± 0.03 | 0.80 ± 0.01, Mouse20, 0.50 ± 0.04 | 0.62 ± 0.03, **Fig. 3b, Supp. Fig. 1c**,**d**).

We further characterized VQ-MAP’s robustness to pose formats by systematically probing the influence of keypoint cardinality and positioning. For keypoint cardinality, we dropped keypoints from the original pose format either at random or following structured patterns that reflect common lab annotation schemes (e.g., one fewer keypoint on each limb, ‘Structural Ablation’, **Fig. 3c**). For keypoint positioning, we applied minor shifts to keypoints by averaging keypoints physically connected along body segments (‘SynAvg’, **Fig. 3c, Methods**). Any effects of these modifications can be directly compared frame-to-frame to the original unmodified data. We compared VQ-MAP to a baseline VQ-VAE model lacking attention modules for keypoint-independent processing and cross-keypoint reasoning after training each model only on unperturbed RatDataset23 data. After training, we assessed whether the models could reconstruct unmodified RatDataset23 pose sequences given perturbed inputs. VQ-MAP was far more robust to changes in keypoint cardinality and positioning over a wide range of perturbation magnitudes and exhibited stable reconstruction performance even when 50% of keypoints were dropped from input poses (**Fig. 3d**). Of the perturbation set, SynAvg was the most disruptive to the baseline model, but VQ-MAP was nearly unaffected (**Fig. 3e**). VQ-MAP also more consistently mapped pose sequences with perturbations to the same behavioral motifs as their unperturbed counterparts (**Supp. Fig. 4**).

**Figure 4:**
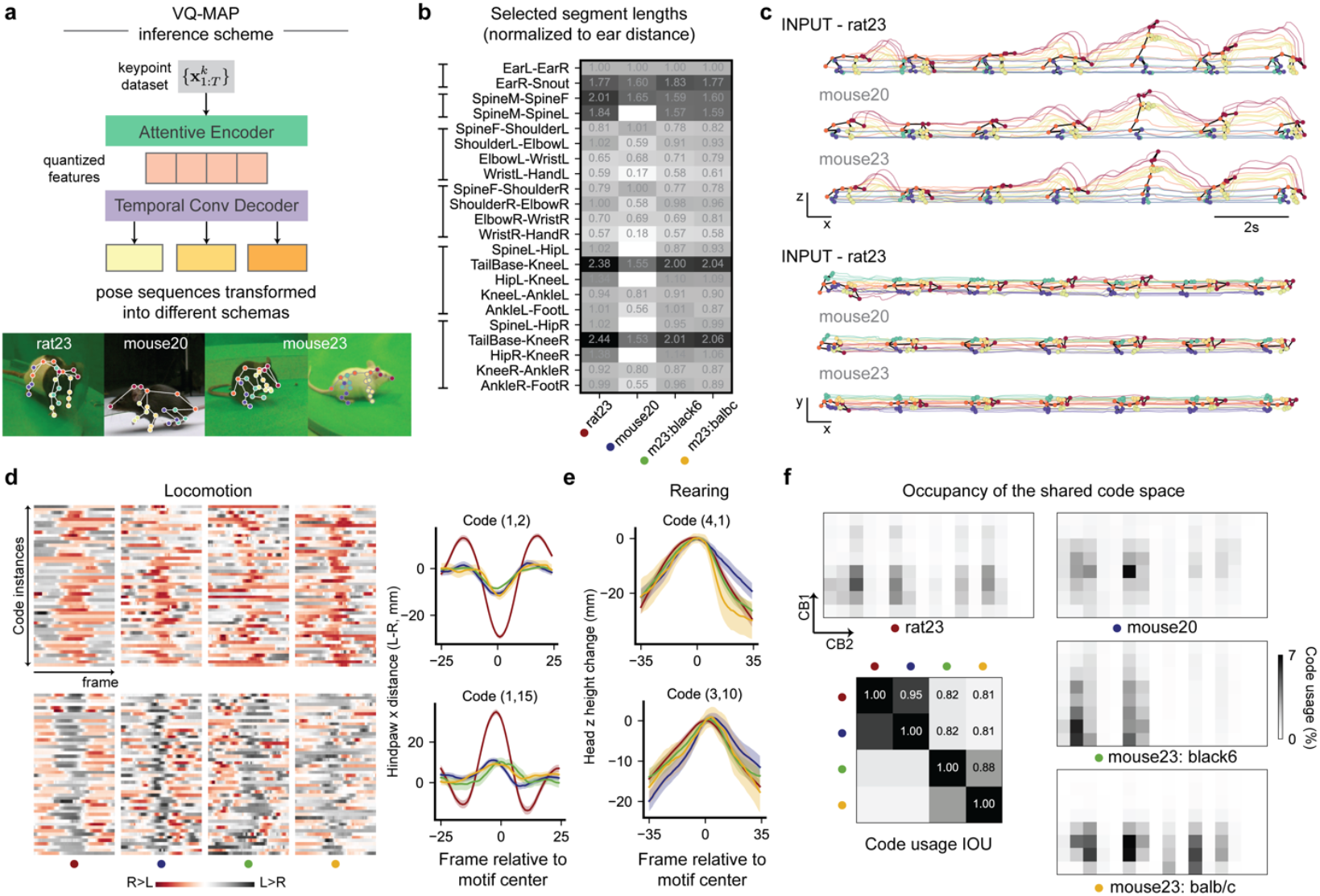
VQ-MAP supports comparative behavioral analyses across species. **a**, Top, illustration depicting how a trained VQ-MAP transforms pose trajectories into different 3D pose schemas by switching the final decoder layers. Bottom, frames from rat and mouse datasets used to train the multi-species VQ-MAP model. **b**, Mean body segment lengths for the different rodent datasets, normalized to the mean inter-ear distance for each dataset. Grayscale levels reflect the values displayed in each cell. **c**, Examples of rat23 pose trajectories converted into mouse20 and mouse23 poses. **d**, Two VQ-MAP motifs tuned to rat and mouse locomotion behaviors. Left, heatmaps showing hindlimb movements (left vs. right limb A-P distances) in n = 50 instances aligned to the same VQ code, top row code (1,2), bottom row code (1,15). Right, mean hindlimb A-P distances aligned to the indicated VQ codes, shaded regions represent 95% CIs. Trace colors and symbols beneath heatmaps indicate different datasets as in **b. e**, Head elevation changes in the z axis aligned to the same VQ codes across different datasets, for codes tuned to rearing. Colors as in **b** and **d. f**, Comparison of VQ-MAP code usage for each dataset. Bottom left, intersection over union (IOU) scores of code usage across different datasets.

We quantified perturbation robustness by training supervised classifiers to recognize MM behavioral labels. When dropping 8 or 13 keypoints from pose inputs, after training only on unperturbed RatDataset23, VQ-MAP maintained 92.4% and 87.1%, respectively, of its maximum classification performance (**Fig. 3f,g**, classifier ‘QuantizedFeatures’). In contrast, baseline model performance fell to 56.9% and 38.8% under the same perturbation conditions. For SynAvg, VQ-MAP achieved 95.5% of its performance on unperturbed poses versus 22.0% for the baseline model. We also found that using VQ-MAP quantized code representations as inputs for action classification was equivalent to using features from other stages of the VQ-MAP network (**Fig. 3g,h**), illustrating how codes conserve behavioral information that can be used for downstream tasks and cross-dataset comparisons.

### VQ-MAP aligns diverse 2D pose datasets

We also applied VQ-MAP to 2D pose datasets, which are far more ubiquitous in the community and include a diversity of pose formats and keypoint cardinalities (cf. **Fig 1a**). To test whether VQ-MAP could bridge common 2D pose format differences, we fit a single VQ-MAP model to three mouse datasets combined: CalMS21^49^ (7 keypoints), KPMS-2D^26^ (8 keypoints), MouseDataset15-2D (15-keypoint subset from top-down 2D projection of 3D mouse poses^42^). While CalMS21 and KPMS-2D have similar cardinalities, their keypoint positioning is markedly different, with KPMS-2D tracking keypoints more densely along the spine rather than on the hindlimbs (**Fig. 3i, Methods, Supp. Table 1**).

We observed no sharp segregation regarding code usage across datasets in the joint behavioral embedding space and identified a set of rearing and locomotion motifs that were shared between the three datasets (**Supp. Fig. 5**). These shared motifs were expressed with similar usage statistics compared to rearing and locomotion motifs identified in VQ-MAP models independently fit to each dataset, suggesting that VQ-MAP accurately collated motifs across different 2D pose formats (**Fig. 3i, Supp. Fig. 6c**,**d**). The CalMS21 dataset included human annotated social behavior labels of the paired mice (attack, mount, investigate, other). Comparing the unsupervised VQ-MAP codes with these social labels (resident mice only), we observed codes that were specifically attuned to each social behavioral category, and more importantly, consistent when trained on CalMS21 alone and when over the combined datasets (**Fig. 3j, left**). We validated VQ-MAP performance on the CalMS21 benchmark dataset, where we found VQ-MAP codes aligned with annotated CalMS21 behavioral categories, both when training VQ-MAP alone and when training over the combined datasets (**Fig. 3j, right**).

**Figure 5:**
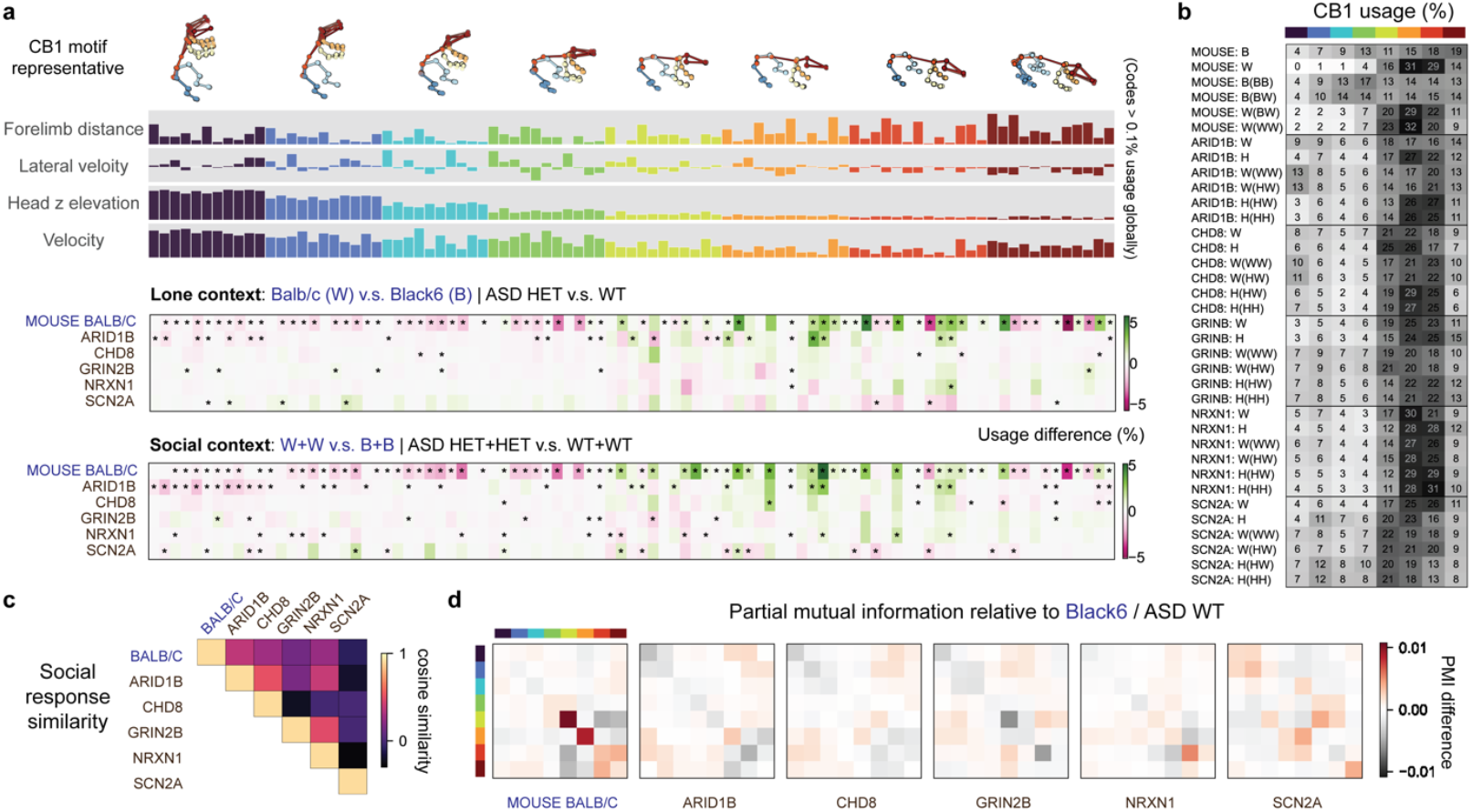
VQ-MAP links between strains and species in rodent autism models. **a**, VQ-MAP code usage differences across mouse strains and five rat models of autism (ASD). Top, representative motifs for each CB1 code and the corresponding kinematic attributes derived from decoded kinematics. VQ-MAP codes were sorted in descending similarity relative to the high rear CB1 motif. For visual clarity, only codes with appreciable use (>0.1%) are shown. Middle and bottom, behavioral differences between heterozygous knockout animals (HET) and wildtype littermates (WT) in lone and social contexts, respectively. BALB/c mice (W) were compared against C57BL/6 (B). **b**, Comparison of VQ-MAP CB1 code usages across species, strains and pairing conditions. Values in each cell represent the percentages of time of each CB1 code expressed in the indicated condition (that is, each row). **c**, Heatmaps showing similarity of behavioral differences (HET vs. WT, W vs. B) in social contexts. Colorbar shows cosine similarity. **d**, Heatmaps showing differences in mean partial mutual information (PMI, colorbar) over CB1 codes for W+W vs. B+B and HET+HET vs. WT+WT.

**Figure 6:**
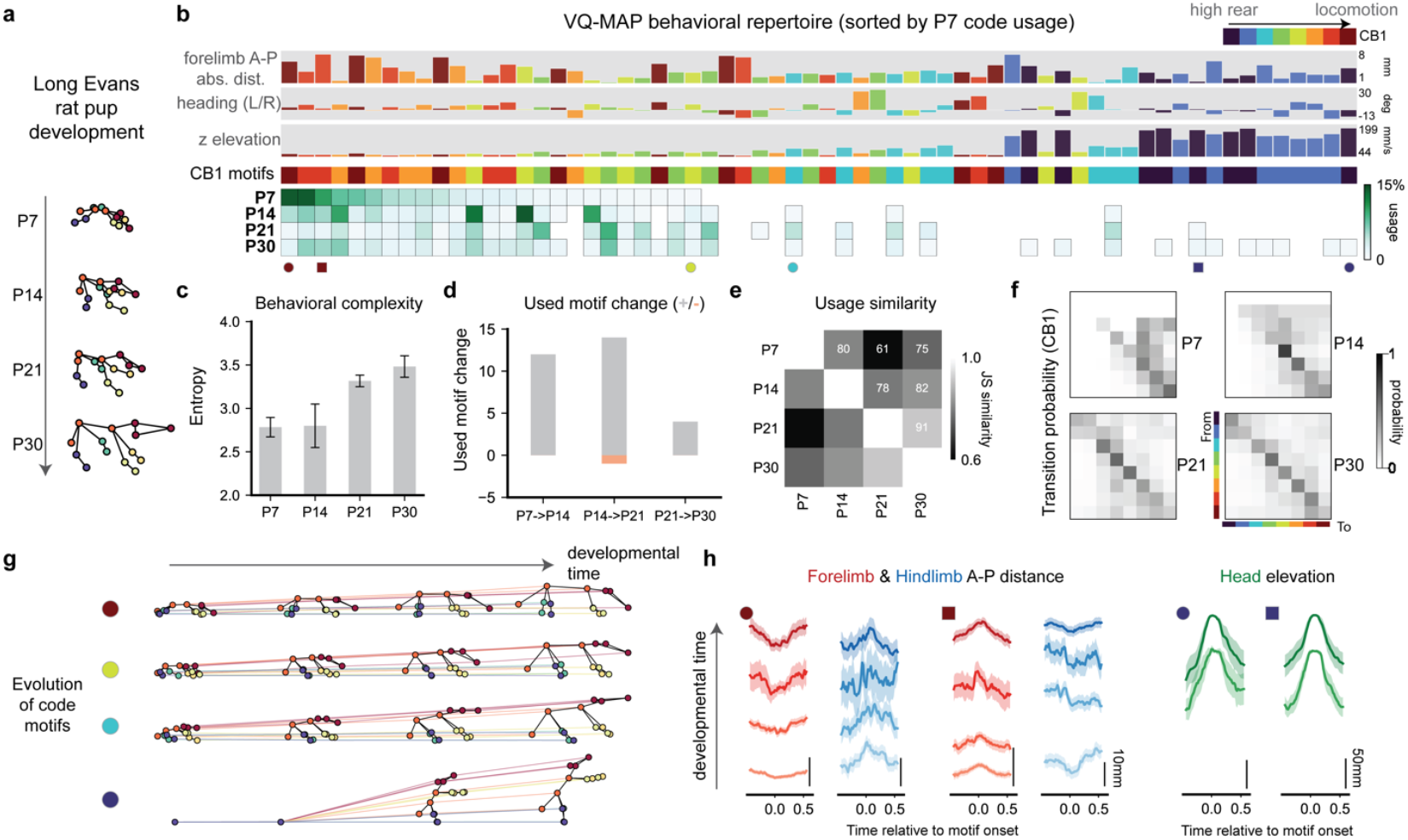
VQ-MAP profiles rat behavioral development. **a**, Example rat postures across four developmental stages, postnatal day 7 (P7), P14, P21, and P30. **b**, VQ-MAP code space usage across all developmental timepoints. Top, VQ-MAP motifs sorted by their usage frequency in P7, together with their basic kinematic compositions. Bottom, code usage across all developmental timepoints. Only codes with usage > 0.5% are displayed. The symbols at the bottom of the heatmap indicate the specific codes highlighted in **g** and **h. c**, Bar plots of behavioral complexity over developmental time, quantified as the entropy of code space usage across codes, mean ± 95% CIs. **d**, The number of new motifs that appeared (gray) or disappeared (orange) between consecutive developmental timepoints. **e**, Heatmap of pairwise behavioral similarity, calculated as the percentage of used codes shared by different developmental timepoints. **f**, Left, transition probability matrices between different CB1 codes for each developmental timepoint. The colors in the P30 plot match those in **b** and denote the type of CB1 code and how these codes are arranged in rows and columns. **g**, Evolution of median motif profiles over developmental timepoints for the codes indicated by symbols in **b. h**, Evolution of limb and head movements across developmental timepoints for the codes indicated by symbols in **b**. Traces are mean ± 95% CIs (shaded areas).

In addition to flexibly supporting different pose formats, VQ-MAP is faster than existing motif identification approaches and readily scales to large datasets with its fully GPU-parallelizable architecture. Compared to KPMS, VQ-MAP shows > 500x speedups during inference on average for every n = 90000 frames (**Supp. Fig. 7**), thus enabling analysis of large-scale, heterogeneous behavioral datasets to support new modes of neurobehavioral discovery.

### VQ-MAP enables comparative behavioral analysis across species

New methods for quantitative behavioral comparison and studies across animal species would shed light on the function and evolution of neurobehavioral systems. To test whether VQ-MAP’s flexibility towards pose format could enable cross-species analyses in real, heterogeneous datasets within the shared behavioral code space, we fit a VQ-MAP model to a combination of RatDataset23, MouseDataset20 and MouseDataset23, where the last contained recordings of two common mouse strains C57BL/6 and BALB/c (**Fig. 4a, Supp. Table 1**). While RatDataset23 and MouseDataset23 were acquired using the same gross 23-keypoint pose schema, segment lengths differed between the datasets owing to natural differences between rat and mouse body plans. Their differences relative to MouseDataset20, which used a separate pose schema, were even more pronounced (**Fig. 4b**).

Once trained, switching the final output layer of the decoder produced pose sequences converted from one species to another, suggesting VQ-MAP learned overlapping codebooks across species and pose schemas (**Fig. 4c**). When we inspected pose trajectory instances mapped to the same VQ codes, we found motifs capturing consistent movement types across species and datasets, which bore substantial similarity with those found in single-dataset VQ-MAP models (cf., **Fig. 2**). For example, codes associated with rodent locomotion exhibited common limb periods and phases (**Fig. 4d**), and rearing codes shared head elevation kinematics across datasets (**Fig. 4e**; cf. **Fig. 2c**).

We observed considerable overlap in the codes expressed by animals in these datasets (intersection over union, IOU, of frequently used codes ranged from 0.81 to 0.95 between datasets, **Fig. 4f**). In contrast, overlap was minimal when we co-embedded RatDataset23 with shuffled mouse datasets (0.14 IOU, **Supp. Fig. 6e**). Motifs identified in this joint model were also consistent with those identified when models were fit to each dataset individually (**Supp. Fig. 6a**,**b**), further supporting the fidelity of dataset-aligned representations.

While the set of used codes was similar across datasets, the usage frequencies of these codes varied by species, strain, and recording environment (**Fig. 4f**). C57BL/6 mice in both MouseDataset20 and MouseDataset23 tended to rear with both hands on the arena wall and body tilted, as well as groom while in a crouching position. However, animals in MouseDataset20 exhibited enriched expression of rearing subtypes characterized by full-body extension, potentially because these mice were recorded in a >6-fold smaller arena than MouseDataset23^50^. The ability to distinguish these subtypes, which captured how behavioral patterns could be perturbed under different experimental conditions yet are expressed primarily in a dataset with a different pose schema, highlights the advantage of VQ-MAP’s joint embedding approach. In these open field recordings, rat motif expression bore many similarities to mice, but rats exhibited a greater diversity of postures within behavioral categories, for example rats often lay prone on the ground or rested on their forelimbs, motifs that were rare in mice.

### VQ-MAP proposes links between strains and species in rodent autism models

An open question in neuropsychiatric and neurological disease research is whether behavioral phenotypes in animal disease models are conserved across species. By facilitating quantitative cross-species comparisons, VQ-MAP offers a new way to probe for such parallels. Here, we used VQ-MAP to investigate mouse and rat models of autism. We applied the previous VQ-MAP model jointly trained with rat and mouse kinematics to a RatAutismDataset of five genetic models of autism (loss of function knockouts of *ARID1B, CHD8, GRIN2B, NRXN1, SCN2A*) and their wildtype littermates (WT), together with the full set of MouseDataset23 (BALB/c, C57BL/6), recorded in both lone and dyadic social contexts (89.48 hours lone and 170.91 hours social from n = 8, 8, 6, 8, 6 *ARID1B, CHD8, GRIN2B, NRXN1, SCN2A* rats; 5.32 hours lone and 17.31 hours social from n = 8 BALB/c and n = 8 C57BL/6 mice, **Supp. Table 1**). BALB/c mice are known to have low sociability^51,52^as a murine model of autism model. However, whether this murine phenotype parallels those in other species and strains is unknown.

To validate VQ-MAP phenotyping, we first compared code space usage patterns within species to previous reports employing MM (**Fig. 5a**). VQ-MAP recapitulated previous observations of lone and social behavioral phenotypes across these rat models of autism, for example, with a social partner present, the decrease in locomotion in *ARID1B* and *CHD8* heterozygous knockouts (HETs; 1.32- and 1.47-fold decrease relative to WT littermates, respectively) and the increased occurrences of rearing behaviors in *SCN2A* HETs (1.22-fold, **Fig. 5a, bottom, Supp. Fig. 8**). The consistent, pronounced behavioral differences between BALB/c and C57BL/6 mouse strains, specifically, the dominant expression of near idle, slow-speed exploration behaviors and much lower rearing activities in BALB/c, were also recovered using the same VQ repertoire (**Fig. 5a,b**).

Using the shared VQ-MAP code repertoire, we compared the behavior of rat HET knockouts and their WT littermates to both mouse strains. We found that for three of the rat KOs (*ARID1B, CHD8, GRIN2B*), BALB/c behavior was most similar to HETs and C57BL/6 to WT littermates, but for *SCN2A* this pattern was flipped, with HETs most similar to C57BL/6 and WTs to BALB/c (**Fig. 5b, Supp. Fig. 8b**). These results resemble a divide reported previously in rats, wherein *SCN2A* and *NRXN1* KOs differed from other rat autism models by displaying increased rates of close social contact^42^.

Of the five rat KOs tested, we found that BALB/c phenotypes were most similar to *ARID1B KOs*, (**Fig. 5c, Supp. Fig. 8**) in terms of the overall distribution of behavioral shifts relative to baselines (C57BL/6 and WT littermates), although not all behavioral effects were conserved. The similarities were driven primarily by shared, significant decreases in the expression of a wide range of vigorous rearing motifs and significant increases in intermediate-speed locomotion and exploration behaviors. We also examined behavioral synchrony between interacting animals and found VQ-MAP CB1 codes recapitulated previous MM results describing changes in synchrony in rat autism KOs (**Fig. 5d**). Patterns in mice differed, with co-occurrences of idle or slow exploration behavior being dominant in pairs of BALB/c relative to C57BL/6.

### VQ-MAP enables novel descriptions and interpretation of behavioral data

We next tested whether VQ-MAP supported quantitative evaluation of the developing rat behavioral repertoire. As body maturation naturally produces differences in pose schema, previous studies built separate behavioral identification models for each animal age and used *post hoc* kinematic similarity metrics to relate behaviors across developmental time^10^. A weakness of this disjoint approach is that all behaviors at a given age are necessarily treated as distinct from the behaviors observed at other timepoints, and thus one cannot reason about whether specific behaviors may be conserved over development.

We revisited the problem using VQ-MAP, fitting a single model to a published dataset^10^ of n = 6 developing Long-Evans rats, recorded at postnatal days (P) 7, 14, 21, and 30 (**Fig. 6a, Methods**). Directly from the shared latent space learned by VQ-MAP, we identified a progressive expansion of rat behavioral repertoire and the emergence of new behaviors over developmental stages (**Fig. 6b, Supp. Fig. 9a**). Consistent with previous reports, behavioral diversity increased over time (**Fig. 6c**), and most new behaviors emerged during the earliest stages (P7→P14, P14→P21, **Fig. 6d**). Animals only started to execute rearing-related behaviors after p7, after which we observed an increasing number of rearing subtypes and maximum rearing amplitude, mostly reflected in the first 3 CB1 codes (‘z elevation’, **Fig. 6b**).

The changes in the rat’s behavioral repertoire were also reflected in transitional probabilities (**Fig. 6f**).

In the VQ-MAP formulation, motifs from the very early developmental timepoints did not disappear from the shared repertoire; instead, via the decoder output layers, they were transformed into larger physical frames with conserved body coordination patterns (**Fig. 6g**). For example, the two most frequent P7 locomotion motifs captured specific limb movement patterns that persisted as body profiles evolved (**Fig. 6g**). Different from previous reports, this revealed an increase in behavioral similarity between P7 and P30, reflecting the tendency of rats in these recordings to locomote (**Fig. 6e**). VQ-MAP thus opens the door to more precise investigations of behavioral development via the shared behavioral encodings.

## Discussion

Pose analysis has become a cornerstone of animal behavior studies, but the inability to relate measurements across experiments, labs, and species has limited its impact. Here, we present how VQ-MAP addresses this issue to enable flexible, joint processing and co-analysis of heterogeneous datasets. Fit to individual datasets, VQ-MAP recapitulates the performance and major findings of the previous studies while offering orders of magnitude speedups in data processing and avoiding the need for *post hoc* hierarchical annotation. Applied across heterogeneous datasets, VQ-MAP seamlessly combines pose formats, including across species, discovering shared patterns of behavioral expression. VQ-MAP thus supports behavioral data standardization and new comparative studies. We have made VQ-MAP an open-sourced codebase (https://github.com/tqxli/vqmap) and have built a graphical interface that facilitates data embedding, visualization and annotation of discovered motifs (**Supp. Fig. 10**).

VQ-MAP’s performance and flexibility stem from features of its network architecture. As VQ-MAP uses attention to combine dynamics from different keypoints, it identifies behavioral representations within inputs of arbitrary size and ordering. VQ-MAP’s quantized latent space also offers distinct advantages. Quantization enables encapsulation of the entire discrete motif identification process in a neural network, permitting facile adoption of network components (attention, hierarchical latent codes, readout adaptors) and enhancement of computational efficiency and scalability via GPUs. Quantized networks are also more memory efficient and easier scaled to large datasets than their continuous network counterparts (standard VAEs), as quantization reduces high-dimensional latent representations to a single-integer code index. VQ-MAP quantization means that discrete behavioral motifs are themselves directly decodable model variables representing specific kinematic profiles. VQ-MAP motifs are not secondary clustering summaries or state variables queried by time-consuming probabilistic iterative sampling. These direct motif representations improve interpretability, training times, and inference speeds.

There is a growing consensus that data collection and analysis pipelines should be standardized to support common behavioral definitions and shared community repositories^53,54^. While we believe it is imperative that the community establish common standards, we expect standardization to be a protracted process. There are thousands of studies that have collected, and likely thousands more that will collect, pose data in idiosyncratic, unstandardized formats. VQ-MAP offers a way to harmonize and link these heterogeneous datasets, a backwards compatible solution supporting community integration before common standards are adopted.

Our results show that VQ-MAP enables analyses over 2D and 3D poses from a variety of pose tracking techniques, but testing over a larger collection of datasets with greater diversity will be important to better understand VQ-MAP’s capabilities and failure modes. For example, it will be valuable to assess how VQ-MAP performs given substantial, non-overlapping behavioral diversity, such as recordings from a complex home cage environment or from disease models that unpredictably alter body kinematics. In such cases, it may prove beneficial for VQ-MAP to explicitly separate shared versus dataset-specific features in separate codecooks^55,56^.

Other limitations of our approach warrant further investigation. VQ-MAP currently uses a fixed codebook timescale. While we chose this timescale (320 ms for rat motif identification) to approximate the typical length of a human-identifiable behavioral motif, future versions of VQ-MAP that flexibly support a range of timescales may better capture motifs that naturally vary in duration. Also, except for in our rat development analyses, we did not explore longer-timescale dependencies in motif expression (e.g., behavioral transitions, sequences, and states). Longer timescale behavioral statistics could be derived via *post hoc* analysis on top of VQ-MAP motifs, akin to learning long-term dependencies over tokenized word sequences using transformer or state space models^57–59^. A future version of VQ-MAP may also explicitly model multiple timescales^60^. Ultimately, the ideal timescale for motif identification depends on the specific application, and there are necessary trade-offs between human interpretability and representational expressiveness.

In the future, we envision VQ-MAP being expanded to support multiple data modalities, including vocalizations, physiological proxies for internal state (e.g., heart rate), and neural activity. VQ-MAP’s solution for behavioral data flexibility might also address the challenge of combining neural data across experiments, where neurons vary in number and are not meaningfully orderable, an area where channel-agnostic models have recently shown promise^61–64^. Extending VQ-MAP to multiple data modalities could enable construction of neurobehavioral foundation models supporting a wide range of analyses and knowledge-building in neuroscience and biomedicine.

## Methods

### Datasets

All animal behavioral datasets used in experiments came from publicly accessible repositories. Dataset details and animal and recording sample sizes are provided in **Supp. Table 1**. For details regarding dataset acquisition and processing, readers should consult the original papers.

### Preprocessing the keypoint pose datasets

For all approaches (VQ-MAP, MotionMapper, Keypoint-MoSeq), we applied the same preprocessing protocol to pose datasets prior to training or evaluation. In each frame, keypoints were transformed into egocentric coordinates, with the middle of the animal’s spine at the origin and the animal’s heading direction facing +x in the xy plane. To determine heading direction, we selected sets of keypoints located at the animal’s anterior and posterior body region for each keypoint schema and computed the direction out of their averaged positions. These sets differed depending on which keypoints were defined in a dataset. Taking RatDataset23 as an example, its anterior keypoints are specified as *EarL, EarR* and the posterior keypoint as *TailBase* (see **Supp. Table 1** for the remaining datasets). Within each dataset, all keypoint coordinates were scaled by a dataset-specific constant to reduce the influence of body size differences across datasets.

For VQ-MAP processing, the keypoint trajectories were segmented into fixed-length sequences of 128 frames and a stride length of 128 frames (that is, sequences did not overlap in time). For each pose schema, keypoints were grouped into 6 general body regions: *Head, Trunk, LeftForelimb, RightForelimb, LeftHindlimb, RightHindlimb* (**Supp. Table 1**).

### VQ-MAP architecture and implementation

#### Nomenclature and mathematical setup

We denote inputs to VQ-MAP as a set of keypoint-based behavioral measurements 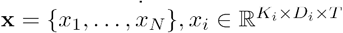 spanning for *T* timepoints, where *N* is the number of datasets with distinct keypoint schemas, *D*_*i*_ = 2,3 is the coordinate dimensionality and *K*_*i*_ is the keypoint cardinality (number of keypoints) associated with the i-th pose dataset. For the heterogeneous data inputs, VQ-MAP aims at learning a shared, discrete latent space ‡ representing stereotyped movement primitives, treating different pose schemas as different projections of the same behavioral dynamics. For an arbitrary model trained on keypoint data with cardinality of *K*_*i*_, which provides a trainable approximation of the posterior distribution of hidden behavioral states **z** conditioned on data inputs **x**_*i*_, it is desirable for it to also allow mapping of (1) a reduced set of keypoints x ^′^ ⊆ x with sufficient information; (2) data with denser keypoints and thus larger cardinality, and (3) measurements **x**_*j*_ collected with novel keypoint placement schemas. The measure of success is the degree to which we can retrieve the same behavioral motifs, both quantitatively when aligned to an existing behavioral repertoire and qualitatively as recognized by humans. We now introduce different components in the unsupervised mapping algorithm in order.

#### Channel-invariant attentive encoder

In the encoder ℰ (·), the keypoint trajectories [*K*_*i*_, *D*_*i*_, *T*] are processed by alternating series of *N*_*enc*_ blocks of (1) temporal convolutional network (TCN) layers that operate independently over each keypoint and downsample the time series by a stride factor *τ* (default *τ* = 2), and (2) an attention-based module that captures cross-keypoint interactions while permitting input cardinality and permutation invariance. The attention module is implemented based on Induced Set Attention (ISA) proposed in Set Transformer^40^. Our attention module performs cross-attention between the input keypoint features 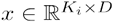 with a learnable query array of embedding vectors ℐ ∈ ℝ ^*M* × *D*^ (“inducing set”) with smaller cardinality than the inputs *M* < < *K*_*i*_.

We use two subsequent standard blocks of CrossAttention(Query, Key, Value) where

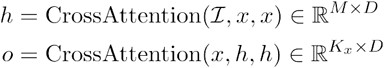

which first transforms the inputs to a smaller cardinality by attending to the inducing query array and later projects to the original cardinality *K*_*x*_. Intuitively, the inducing set groups and aggregates features from a variable number of channels via pairwise similarity. To remove dataset-specific information (i.e., keypoint cardinality) from the encoder architecture, we omit the second CrossAttention block and directly output *h* ∈ ℝ ^*M* × *D*^ after the final TCN layers. The timescale 𝒯 captured by each discrete latent code is therefore determined by the number of encoder processing blocks and their corresponding temporal stride in the TCN layers. For all behavioral analysis experiments comparing code usage, we used *N*_*enc*_ = 4 blocks each of stride. For the action classification transfer experiments, we used *N*_*enc*_ = 4 blocks of stride.

#### Body region encoding

While removing keypoint identity entirely would be most flexible, in practice we found it helpful to retain information about keypoints at the body region level, given how sparsely distributed keypoints are across the body. To achieve this, while maximizing generalizability towards different pose formats, we adopted a positional encoding scheme that incorporated the gross body part location for each keypoint. Prior to training, we grouped keypoints within each pose schema into one of 6 body regions (*Head, Trunk, Left Forelimb, Right Forelimb, Left Hindlimb, Right Hindlimb;* **Supp. Table 1**). During training, we injected corresponding gross body location information via learnable encodings (jointly updated with the VQ model) into each keypoint dimension after pose sequences were processed by the first TCN block. For inputs 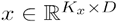, BodyPartEncoding(*x*) = *x* + *A*^*T*^*W*_*BP*_*x* where *P* denotes the number of general body parts, 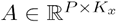 is an indicator matrix used to assign encoding vectors from *W*_*BP*_ ∈ ℝ ^*P* × *D*^ to the corresponding channels.

#### Differentiable vector quantization

The channel-invariant encoder ℰ (·) provides a trainable approximation of the posterior distribution of latent codes 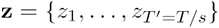 conditioned on **x**, a sequence of latent variables whose length depends on the overall temporal downsampling factor *s*. The prior distribution is chosen as uniform over a dictionary of embedding vectors {***e***_*i*_ ∈ ℝ ^*D*^}, *i* ∈{1, …,*K*} which collectively forms a discrete latent space ℰ ∈ ℝ ^*K* × *D*^ where *D* is the dimensionality of each embedding vector. The posterior is implemented as a one-hot categorical distribution *q*_*ϕ*_ (***z*** = *k* | ***x***) = 1 for *k* = argmin_*i*_ ‖ ***z*** − ***e***_*i*_ ‖_2_ (otherwise 0), which retrieves and quantizes the input features with the nearest neighbor drawn from the codebook. The vector quantized representations therefore equal ***z***_*q*_ (*x*) = *VQ* (*z*) = ***e***_*k*_ where *k* = argmin_*i*_ ‖ ***z*** − ***e***_*i*_ ‖_2_. Given that is *p*(***z***) uniformly distributed over embedding vectors, the associated KL divergence (if viewed as a standard variational autoencoder) 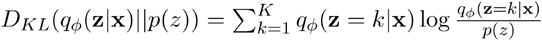 reduces to a constant of log *K* and can be ignored in the training objective.

Given the non-differentiability of nearest-neighbor lookup operation from the embedding space, a straight-through estimate is made by directly copying the gradient from decoder inputs ***z***_*q*_ after quantization to the encoder outputs ***z***(***x***). PyTorch pseudocode for this process is:

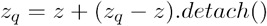

Please refer to the *Optimization objective* on specific implementations to enable this optimization.

To minimize information loss from quantization, we based our vector quantization implementation on residual vector quantization (RVQ)^65^, a variant of the original VQ-VAE implementation^33^ named r. In RVQ, a stack of quantizers is used, where all quantizers following the first quantize the residuals from the last level. We used 2 levels in each residual vector quantizer and only used the top-level codebooks for analysis.

#### Multi-codebook product quantization

We deployed a parallel set of residual vector quantizers with codebook size *N*_1_, …, *N*_*C*_ (top-level codebooks only for RVQ), which independently quantize equal-sized partitions of. This “product quantization” strategy^47^ improves code diversity from 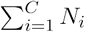 to 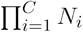 while maintaining a sufficiently small total number of distinct codes. The posterior can be then rewritten as *q*_*ϕ*_ (***z*** |***x***) = *q*_*ϕ*_ (***z***^1^, …, …, ***z***^*C*^|***x***).

To encourage disentanglement of generative factors captured in each codebook, we applied behavior-invariant transformations to the training datasets, including left-right pose flipping and keypoint dropouts (at most 3 keypoints were dropped from each body region, with each keypoint in the pose dropped with a fixed probability). During training, we introduced an “assignment loss” term in the learning objective, to encourage CB1 encodings of transformed versions to be similar.

#### Multi-head decoder

The decoder network 𝒫 (·) is fully convolutional, consisting only of TCN layers that gradually upsample the latent features 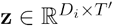 by reversing encoder network striding. Depending on training dataset formats, the end of the decoder is equipped with different output heads that map hidden features [*T,D*_*h*_] to specific pose schemas [*T,K*_*i*_ * *D*_*i*_] for computing the reconstruction loss.

#### Optimization objective

The overall learning objective is given by

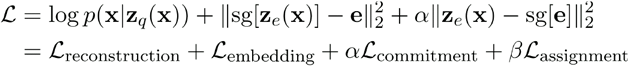

where sg represents the stopgradient operator with zero gradients.

The first term represents the negative log-likelihood of the reconstruction, as in a typical VAE formulation. However, the straight-through gradient estimate prevents this reconstruction term from establishing gradients on the codebook itself. The embedding loss is an L2 loss geared towards the codebook, encouraging learning of code embedding vectors that are similar to encoder features. In practice, we achieved better performance when implementing the embedding loss as an exponential moving average (EMA) of encoder outputs. The commitment loss constrains the embedding space volume and acts as a regularizer^33^. As described above in *Multi-codebook product quantization*, the assignment loss promotes hierarchical encoding of behavior.

To summarize, in the learning objective, the first reconstruction term optimizes the encoder and decoder; the second embedding term (implemented with EMA) optimizes the embedding vectors in the codebook; the last commitment term only optimizes the encoder. The commitment term is scaled by a hyperparameter *α* in the final objective. We used *α* = 0.002 following existing papers for human motion quantization ^37,38^

#### VQ-MAP model training

We trained all models using a single 24GB memory GPU (RTX 3090 or A5000). We used an AdamW optimizer with *β*_1_ = 0.9, *β*_2_ = 0.99 and weight decay=0.0 and used a cosine annealing schedule with an initial learning rate of 0.0001 and cycle period of 50, for a total of 450 epochs. Left/right flipping transformations were applied to every input, with equal probability of left vs. right, and keypoints were dropped with a probability of 0.2, with at most 3 keypoints dropped per body region.

### Model evaluation

#### Visualizing VQ-MAP behavioral motifs

We used two strategies to visualize the pose trajectories associated with each VQ-MAP code motif: (1) we fed sequences comprising single, repeated latent codes into the VQ-MAP decoder, which directly outputted their associated motif pose trajectories (e.g., **Fig. 2a**), (2) we embedded a dataset and computed the median trajectories out of all instances mapped to the same VQ code. For (1), the pose trajectories are of fixed duration as determined by the encoder temporal strides, e.g., 16 timepoints for a 4-layered encoder of stride 2. For (2), we additionally padded around each motif instance into 1s sequences, e.g., 16 + 2 * 17 = 50 timepoints for a recording of 50 fps.

#### Inference speed benchmarking

We measured the time it took for trained VQ-MAP and KPMS models to run inference over a 30-minute RatDataset23 recording (**Supp. Fig. 7a**). The inference time was measured on a single RTX3090 GPU, with batch size set to the maximum permitted by GPU memory (24 GB). For KPMS, the fitting time was measured for 500 iterations (the KPMS default) and 100 iterations. Additionally, we measured the single-batch memory consumption (MB) of VQ-MAP as a function of input keypoints (**Supp. Fig. 7b**). We synthesized data batches of shape [batch size = 1, T = 128, K = k, D = 3], as the number of keypoints ranges between 10 and 50, and recorded the maximum GPU memory allocation during forward and backward pass using ‘torch.profiler’. We reported the memory consumption averaged over 10 iterations for each data batch.

#### Baseline model implementation

We compared VQ-MAP with a baseline VQ-VAE model that replaced the keypoint cardinality-invariant encoder with a channel-mixing encoder (**Fig. 3**). For channel-mixing, we used an encoder architecture that was symmetric to the decoder, that is, the encoder consisted of different input linear processing layers to accommodate different pose formats (Linear(KxD, 128) for a pose format of K keypoints, D=2 or 3) followed by shared TCN blocks.

#### Cross-dataset motif regression

To assess consistency across different motif mapping results on the same dataset, we compared one motif mapping (MotifMappingA) to the other (MotifMappingB), using linear regression with 10-fold cross-validation (Python sci-kit learn, sklearn.linear_model.LinearRegression). When comparing MotifMappingA with MotifMappingB, we fit linear regression models that fit A to B and vice versa and reported the average train and test R-squared values and Hamming distances between model predictions and the target motifs.

### Comparison to other behavioral analysis frameworks

We compared the behavioral motif identification performance of VQ-MAP to two published unsupervised behavioral mapping methods, Keypoint-MoSeq and MotionMapper.

#### Keypoint-MoSeq (KPMS)

We fit KPMS models using its default parameters, κ = 1*e*4, latent dimension of 16, with 50 iterations of AR-HMM fitting and 500 iterations of full model fitting. For cross-dataset mapping experiments (**Fig. 1b,c**; **Fig. 2h,i, Supp. Fig. 1**), we manually ordered keypoints in the inference dataset to match keypoint order in the training dataset. When a KPMS model was applied to a dataset with a different keypoint cardinality, for the inference dataset we used a common keypoint subset (when inferring over a dataset with higher cardinality) or keypoint duplication (when inferring over a dataset with lower cardinality). For example, when applying a RatDataset10 KPMS model to RatDataset23, we only used only the subset of rat23 keypoints present in rat10 pose configuration. When applying a RatDataset23 KPMS model to RatDataset10, we inserted rat10 keypoints into their corresponding positions within the rat23 pose configuration, filling rat23 keypoints not in rat10 with zeroes. For KPMS co-training with datasets of different cardinality, we report results from two models, one where we reduce pose configurations to the lowest cardinality via subsetting, and one where we increase pose configurations to the highest cardinality via zero padding.

#### MotionMapper (MM)

We used the MM outputs provided with the social-DANNCE dataset^42^ for comparisons on RatDataset23. A detailed description on the MM procedure is provided in Klibaite et al.^42^. Briefly, MM identifies a rich repertoire of behavioral motifs from clustering high-dimensional postural-temporal features, constructed via wavelet decomposition on top eigenpostures, in a lower dimensional t-distributed stochastic neighbor embedding (t-SNE) embedding space. Two human labelers annotated each cluster with a fine-grained behavioral description and assigned a coarse behavioral category out of *idle, sniff/head, groom, scrunched, active crouched, rear, explore, locomotion*.

#### VQ-MAP

By default, for graphs and visualizations we sorted CB1 codes in descending order of similarity to the CB1 code with the highest average head height (relative to the animals’ feet) in the decoded kinematics across all combinations with CB2. We defined similarity as the cosine similarity between code embedding vectors. Readers can refer to **Supp. Fig. 2a** for visualization of a resulting code space after sorting. Because VQ-MAP does not require data structure shape to match between training and inference datasets, when applying VQ-MAP across datasets of different cardinality, no compatibility adjustments to pose configurations were imposed.

### Behavioral analysis

#### Behavioral similarity

To compare two behavioral experiments at the level of overall motif expression patterns, we normalized the raw code space occupancies into probability distributions *P*_1_ for experiment 1 and *P*_2_ for experiment 2 and computed the Jensen-Shannon divergence 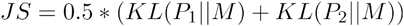, where M = (*P*_1_ + *P*_2_)/2 and KL is the Kullback-Leibler divergence 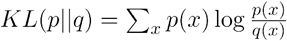 for discrete distributions p and q. The behavioral similarity is defined as 1 - (JS / log(2)), which is bounded between 0 and 1.

### Response similarity

When examining similarity of behavioral changes (e.g., lone vs. social context **Fig. 4c**), we computed cosine similarity between the two feature vectors 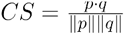.

#### Supervised action classification

We trained action classifiers using RatDataset23 and human-annotated MotionMapper labels (n = 8 action classes). The action classifiers were 3-layer unidirectional LSTM networks with a Linear layer for logit outputs, trained using a standard cross entropy objective. We tested four input data scenarios: (1) unmodified rat23 pose sequences of 128 frames in length, (2) intermediate feature representations of these sequences from a pretrained VQ-MAP model encoder prior to quantization, (3) intermediate feature representations of these sequences from a pretrained VQ-MAP model encoder after quantization, and (4) these sequences passed through and decoded by VQ-MAP but before the final linear readout layer. Because the VQ-MAP encoder features (scenarios 2 and 3) are downsampled relative to the input, for these scenarios we y interpolated so that classifier inputs matched the length of the original pose sequences.All action classifiers are trained for 35 epochs using an Adam optimizer and a constant learning rate of 0.001.

### Statistical analysis

To compare different experimental groups in **Fig. 5a**, for a specific VQ-MAP code, we fit linear mixed effects model (’statsmodels.formula.api.mixedlm’) to its usages across different recording sessions (‘observations’), with a fixed effect for the experimental condition (i.e., ASD KOs vs. WTs, BALB/c vs. C57BL/6 in lone or social context) and a random effect for subject identities. We applied Benjamini-Hochberg false discovery rate correction (’statsmodels.stats.multitest.fdrcorrection’) to the obtained p values (n = 91, global usage > 0.1%).

### Software

To aid reproducibility and help researchers analyze custom behavioral datasets, we have made VQ-MAP freely available as a Python package at https://github.com/tqxli/vqmap, which can be conveniently installed into a Conda environment. The codebase is licensed under a Creative Commons Attribution 4.0 International License. The codebase includes:

- ’configs’: a configuration system supported by Hydra that allows users to specify experiment parameters in a hierarchical, modular fashion via YAML files and maximizes reproducibility.
- ’core/train.py’ and ‘core/inference.py’: main entry points for interacting with the source codes.
- ’dataset’: dataset class definitions and utility functions for egocentrically align the pose datasets.
- ’model’: implementation of VQ-MAP networks solely with PyTorch.
- ’gui/main.py’: a PyQt5-based graphical interface for the users to annotate and analyze data using a pretrained model.
- ’demo’: tutorial Python scripts and Jupyter notebooks.

Users should refer to the GitHub ‘README.md’ for additional instructions on codebase usage.

## Data and code availability

The code for dataset preparation, training and evaluation of VQ-MAP models is made available at https://github.com/tqxli/vqmap. We have also provided an interactive graphical interface implementation that allows users to visualize and annotate the VQ-MAP motifs with finer-grained labels, as well as update and embed different keypoint datasets using pretrained models.

## Acknowledgements

This work was supported by funding from SFARI (BTI Fellowship) to U.K., the National Science Foundation (GRFP and INTERN fellowships) to J.H.W., and the National Institutes of Health (R34DA059506, R34DA059512, and R01GM136972) to T.W.D.

## Author contributions

Conceptualization, T.L., J.H.W., and T.W.D.; Methodology, T.L. and T.W.D.; Software development, T.L.; Validation, T.L. and T.W.D.; Formal analysis, T.L.; Investigation, T.L., J.K., U.K.; Writing – original draft preparation, T.L., U.K., J.H.W, T.W.D.; Writing – review & editing, T.L., U.K., and T.W.D.; Supervision, T.W.D.; Funding acquisition, U.K. and T.W.D.

